# KAT8 facilitates the proliferation of cancer cells through enhancing E7 function in HPV-associated cervical cancer

**DOI:** 10.1101/2024.12.26.629157

**Authors:** Anli Xu, Xiaoming Yang, Junwei Zhao, Shujun Kong, Qing Tang, Xiangzhi Li, Hongmei Qu, Guoyun Wang

**Author notes:** These authors contributed equally to this work (Anli Xu, Xiaoming Yang). Correspondence address.; (G.W.) / (H.Q.) / (X.L.).

## Abstract

Persistent human papillomavirus (HPV) infection serves as the principal etiological factor in cervical cancer, with the oncoprotein E7, encoded by the virus, playing a key role in tumorigenesis. Despite this, targeted therapeutic strategies against E7 remain underexplored. KAT8, a lysine acetyltransferase, significantly contributes to oncogenesis through the regulation of transcription. However, its involvement in cervical cancer remains inadequately characterized. This study employs HPV18-positive HeLa and HPV16-positive SiHa cell lines to investigate how KAT8 modulates E7 expression and function in cervical cancer cells. Upon KAT8 knockdown, a marked reduction in cell viability is observed, alongside a downregulation of E7 expression. This is associated with elevated levels of retinoblastoma protein (pRb) and decreased E2F1 expression, indicating that KAT8 depletion inhibits E7 expression, resulting in E2F1 inactivation and cell cycle arrest. Furthermore, KAT8 is found to directly bind to the promoter regions of the HPV18 LCR, enhancing transcription of the HPV18 E7 gene. This study also demonstrates that KAT8 is essential for the acetylation of E7, and plays a critical role in facilitating the interaction between pRb/E2F1 and E7 in cervical cancer cells. In conclusion, these results highlight KAT8 as a key driver of cervical cancer progression, promoting HPV E7 expression and its associated oncogenic signaling pathways.

## Introduction

Cervical cancer ranks among the most common gynecological malignancies, with squamous cell carcinoma (SCC) and adenocarcinoma (ADC) representing the primary histological subtypes, collectively accounting for 70% and 25% of cases, respectively [1]. The disease originates from normal cervical epithelial cells, progressing through low-grade and high-grade cervical intraepithelial neoplasia (CIN) before advancing to invasive cancer. Persistent infection with high-risk human papillomaviruses (HR-HPVs) is central to its pathogenesis [2]. Among these, HPV types 16 and 18 are implicated in over 80% of cervical cancer cases. Although vaccines targeting oncogenic HPV strains have been developed, they do not eradicate existing infections nor prevent their progression to malignancy [3, 4]. As a result, cervical cancer continues to pose a significant global health threat, underscoring the need for innovative therapeutic strategies for HPV-related cervical cancer.

The HPV oncoproteins E6 and E7 play integral roles in the virus’s life cycle and the transformation of infected cells [5]. Despite lacking intrinsic enzymatic activity, E6 and E7 mediate the degradation of critical tumor suppressors, p53 and pRb, respectively, by recruiting the ubiquitin-proteasome system [4]. Specifically, the interaction between E7 and pRb results in the dissociation of E2F1 from the E2F1/pRb complex, thereby promoting the transcriptional activation of E2F1-regulated genes and advancing the cell cycle into the S phase [6]. In addition, HPV16 E7 directly binds to E2F1, driving E2F1-dependent transcription independently of pRb. Furthermore, E7 influences cellular proliferation through epigenetic modulation, enhancing DNMT1 activity and promoting hypermethylation of tumor suppressor gene promoters [3]. These epigenetic modifications lead to gene silencing, cellular transformation, and tumorigenesis. E7 also alters histone post-translational modifications (PTMs), including the activities of histone acetyltransferases (HATs) and histone lysine demethylases (KDMs), thus reshaping the epigenetic profile of additional oncogenes in cervical cancer cells [7-9]. These findings highlight the therapeutic potential of targeting HAT or KDM inhibitors in HPV-associated cervical cancer [10]. While substantial progress has been made in elucidating the biochemistry of HR-HPV E7, the molecular mechanisms governing its function remain unclear.

KAT8, a member of the MYST family of HATs (also known as MYST1 or MOF), catalyzes the acetylation of Lys16 on histone H4 (H4K16), a modification that promotes gene transcription. Beyond histone modification, KAT8 acetylates various non-histone substrates, influencing their functional properties [11]. This enzyme plays an essential role in gene expression activation, chromosomal stability maintenance, cell cycle regulation, DNA damage repair, and early embryonic development [12-14]. Disruption of KAT8 function has been linked to the development of several diseases, particularly cancers, although the precise mechanisms remain poorly understood. Loss of KAT8 has been shown to exacerbate the genotoxic effects of cisplatin in HeLa cervical cancer cells [15]. Additionally, KAT8 regulates the expression of genes involved in oxidative phosphorylation and contributes to the repair of respiratory defects in HeLa cells [15]. These results suggest a potential role for KAT8 in modulating the biological processes of cervical cancer cells, though the underlying mechanism remains undefined.

This study reveals a significant association between elevated KAT8 expression in cervical cancer and poor patient prognosis. Furthermore, KAT8 enhances the activation of the E7/E2F axis, driving the proliferation and metastasis of cervical cancer cells *in vitro*. The acetylation of HPV16/18 E7 by KAT8 is essential for the functional activity of E7 in HPV-driven oncogenesis. In conclusion, the findings highlight the integral role of KAT8 in the pathogenesis of HPV-induced cervical cancer.

## Material and Methods

### Cell culture and reagents

HEK293T cells, along with human cervical cancer cell lines HeLa (HPV18-positive), Caski (HPV16-positive), and SiHa (HPV16-positive), were sourced from Wuhan Pricella Biotechnology Co. Ltd. The human cervical epithelial cell line HcerEpic was obtained from Sunncell Biotech Co., Ltd. HeLa and SiHa cells were cultured in MEM medium containing 10% fetal bovine serum (Procell, PM150410) and 1% penicillin-streptomycin (Solarbio, P1400). HEK293T cells and HcerEpic cells were grown in DMEM medium, while Caski cells were cultured in RPMI-1640. All cell lines were maintained at 37°C in a humidified incubator with 5% CO_2_. HeLa cells underwent treatment with the KAT8 inhibitor MG149 (MCE, HY-15887) for 24 hours.

### Plasmids and cell transfection

Stable KAT8 overexpression (KAT8-OE) and knockdown (shKAT8) cell lines were established through lentiviral transduction. The lentiviral vectors pLenti-U6-shKAT8-CMV-puro and pLenti-GIII-CMV-Flag-KAT8 were synthesized by Abm (Jiangsu, China). After 24 hours of lentiviral infection, cells underwent selection with puromycin (1 μg/mL) to obtain stable lines. Transfection efficiency was validated via Western blot analysis. The shRNA sequences were as follows: shKAT8-1#: ATGCTGTACAGAAGAACTCATTCAAGAGATGAGTTCTTCTGTACAGCAT; shKAT8-2#: AGACAGTGAAGGATGCTGTATTCAAGAGATACAGCATCCTTCACTGTCT; shNC: GGGTGAACTCACGTCAGAATTCAAGAGATTCTGACGTGAGTTCACCC. Cells were subsequently transfected with plasmids (pLenti-GIII-CMV-HA-HPV16/18-E7) using a transfection reagent (Polyplus, 101000046). Subsequent analyses were performed 48-72 hours post-transfection.

### Cervical cancer specimens

A cohort of 43 patients with cervical squamous cell carcinoma or adenocarcinoma, identified as HPV+/HPV-from Yantai Yuhuangding Hospital from 2022 to 2024, was selected for the study. This cohort included 39 cases of cervical squamous cell carcinoma and 4 cases of cervical adenocarcinoma, with 38 HPV-negative and 5 HPV-positive cases. Inclusion criteria conformed to the established diagnostic standards for cervical cancer in Gynecology and Obstetrics. Exclusion criteria encompassed: the presence of other malignancies; recent chemoradiotherapy, immunosuppressant, or interferon treatments; autoimmune diseases; and incomplete cervical anatomy due to prior partial or total resections. Ethical approval for the study was granted by the Ethics Committee of Yantai Yuhuangding Hospital (approval number: 2021638). Informed consent was obtained from all participants prior to their inclusion, and the study was conducted in strict adherence to ethical principles and regulations.

### Western blot analysis

Cell lysates were obtained by incubating cells with lysis buffer (Beyotime, P0013) containing a 1% protease inhibitor cocktail (MCE, HY-K0021). For protein separation, 30 µg of total protein was loaded onto an SDS-PAGE gel and transferred to PVDF membranes (Millipore, USA). After blocking with 5% BSA, membranes were probed with primary antibodies overnight at 4°C, followed by incubation with HRP-conjugated secondary antibodies. Immunoreactive bands were visualized using the ChemiScope6200 Touch (Shanghai, China). The following primary antibodies were employed: anti-KAT8 (1:1000, abcam, ab200660), anti-β-actin (abclonal, AC038, 1:10000), anti-HPV18 E7 (abcam, ab100953, 1:1000), anti-HPV16 E7 (Invitrogen, 28-0006, 1:1000), anti-E2F1 (Proteintech, 66515-1-Ig, 1:1000), anti-pRB1 (S807) (Cusabio, CSB-RA019386A807phHU, 1:1000), anti-Pan Acetyl-Lysine (abclonal, A1525, 1:1000), anti-HA-Tag (abclonal, AE008, 1:1000), and anti-Flag-Tag (abclonal, AE004, 1:1000).

### Quantitative real-time PCR (RT-qPCR)

RNA was extracted using RNA-easy Isolation Reagent (Vazyme, R701) according to the manufacturer’s instructions. cDNA synthesis was performed with random primers and HiScript II Q RT SuperMix with gDNA wiper (Vazyme, R223). PCR amplification was carried out using ChamQ Universal SYBR qPCR Master Mix (Vazyme, Q711) under the following cycling conditions: initial denaturation at 95°C for 30 seconds, followed by 40 cycles of 95°C for 10 seconds, 60°C for 30 seconds, and 72°C for 30 seconds. Gene expression was quantified in triplicate and normalized to GAPDH levels. Data are presented as the mean of three independent biological replicates. Primer sequences used were: KAT8-forward: TCACTCGCAACCAAAAGCG, KAT8-reverse: GATCGCCTCATGCTCCTTCT, GAPDH-forward: TTCTCTGCGTCGTTGGAGTC, GAPDH-reverse: GGCTGTTG TCATACTTC TCATGG.

### Cell Counting Kit-8 (CCK-8) Assay

Cell viability was evaluated using the Cell Counting Kit-8 (CCK-8) assay (B34304, Bimake). For this, cells were plated in 96-well plates at a density of 1 × 103 cells per well. CCK-8 reagent was added at designated time intervals, and cells were incubated at 37°C for 1 hour. Absorbance was then measured at 450 nm using a Microplate Reader.

### Colony formation assays

For colony formation assays, 600 cells were seeded in 6 cm plates and incubated in complete growth medium for 10-14 days. Colonies were then fixed in methanol for 10 minutes and stained with crystal violet for 15 minutes at room temperature. After washing extensively with water to eliminate excess dye, colony counts were obtained using ImageJ software.

### Transwell assay

Transwell assays were performed using 8-μm pore size transwell inserts (BD Biosciences, USA) placed in 24-well plates, with or without Matrigel (BD Biosciences, USA). In migration assays, 5 × 10□ cells were seeded in the upper chamber in 200 μL of serum-free medium, while for invasion assays, 3 × 10□ cells were used. The lower chamber contained 700 μL of medium supplemented with 10% FBS. After incubation at 37°C for the designated time, cells that migrated to the lower membrane surface were fixed with methanol, stained with 0.1% crystal violet, and subsequently observed and quantified using a light microscope.

### Wound healing assay

Wound healing assays were conducted by plating cells in 6-well plates. Upon reaching 90% confluence, a sterile 200 μL pipette tip was employed to create a scratch wound. Cells were cultured until confluence was re-established, and images were taken at 0, 24, and 48 hours.

### Cell cycle assay

For cell cycle analysis, HeLa and SiHa cells were trypsinized, fixed in 70% ethanol, and stained with propidium iodide (PI). After staining, cells were analyzed via flow cytometry according to the manufacturer’s protocol (Liankebio, CCS01). Data were processed using ModFit software for cell cycle analysis.

### TUNEL assay

Apoptosis in HeLa and SiHa cells was evaluated by immunofluorescence (IF) following the manufacturer’s protocol (KeyGEN C1089). Cells were fixed with 4% paraformaldehyde at room temperature for 30 minutes, washed thrice with PBS, and treated with DNase I reaction solution. Subsequently, TdT enzyme reaction solution, Streptavidin-TRITC labeling solution, and DAPI were applied. Fluorescent images were captured using a Carl Zeiss Observer7 microscope.

### Cell apoptosis assay

Cells were digested with trypsin (without EDTA) and washed with PBS, followed by collection of 1-5 × 10□ cells. Apoptosis was assessed using the Annexin V-FITC apoptosis detection kit (APExBIO K2003), with cells stained by Annexin V-FITC and propidium iodide (PI). Apoptotic cell detection was performed *via* flow cytometry (FACS), according to the manufacturer’s instructions (APExBIO K2003).

### Tissue immunohistochemistry

Tissue samples were deparaffinized in xylene, rehydrated through a graded ethanol series, and subjected to antigen retrieval in citric acid. After blocking with normal serum, slides were incubated overnight at 4°C with an anti-KAT8 primary antibody. Following incubation, the slides were stained using the DAB Kit according to the manufacturer’s protocol (Gene Tech, GK600510). Imaging was conducted with a LEICA DMLB2 microscope at 4× and 20× magnification. IHC scores for cancerous and adjacent cervical tissues were calculated using Image-Pro Plus 6.0. The score was derived using the following formula: (percentage of cells with weak intensity × 1) + (percentage of cells with moderate intensity × 2) + (percentage of cells with strong intensity × 3).

### Coimmunoprecipitation (CoIP)

To assess the interaction between KAT8 and E7, CoIP was conducted by co-transfecting Flag-KAT8 and HA-HPV16/18-E7 overexpression plasmids into HEK293T cells. Cell lysates, prepared in IP lysis buffer (pH 7.4, 0.025 M Tris, 0.15 M NaCl, 0.001 M EDTA, 1% NP40, 5% glycerol), were incubated overnight at 4°C with anti-Flag and anti-HA antibodies, or with IgG as a control. Immune complexes were then incubated with protein A/G magnetic beads for 1 hour at room temperature, followed by washing to remove unbound material. Bound complexes were eluted from the beads using low-pH buffer (Beyotime, P2179S) and analyzed by Western blot.

### Chromatin immunoprecipitation (ChIP) assay

ChIP assay was performed using the SimpleChIP® Plus Enzymatic Chromatin IP Kit (Magnetic Beads) (Cell Signaling Technology, #9005). HeLa cells (4 × 10^4^ cells per IP) were treated with 1% formaldehyde for 20 minutes to crosslink proteins to DNA. Following cross-linking, cells were harvested. Chromatin DNA was digested with micrococcal nuclease and fragmented by sonication to an average size of 150–750 bp.

Protein G magnetic beads were employed to immunoprecipitate chromatin-DNA complexes using KAT8 or IgG antibodies. DNA was purified according to the established protocol. Purified DNA was subsequently analyzed by RT-qPCR. Primer sequences used were: Primer-1#-forward: CGAAATAGGTTGGGCAGCAC; Primer-1#-reverse: TCCCGACCGTTTTCGGTTAC; Primer-2#-forward: TTGGGCACTGCT CCTACATATT; Primer-2#-reverse: AATTGTTGTAGCGCACCTGGA.

### RNA-sequencing (RNA-Seq) and data analysis

RNA samples from control and KAT8-OE HeLa or SiHa cells that passed quality control were sent to OBiO Technology (Shanghai) for library preparation and subsequent sequencing on the Illumina platform. Differentially expressed genes (DEGs) were identified based on a *P* value < 0.05 and |Log2(Fold Change)| > 1. The selected DEGs were subjected to Gene Ontology (GO) and Kyoto Encyclopedia of Genes and Genomes (KEGG) pathway analysis, with GO and KEGG annotations retrieved using the DAVID tool. A significance threshold of *P* value < 0.05 was applied in both GO and KEGG analyses.

### Statistical Analysis

Survival curves for cervical cancer patients were generated using the Kaplan-Meier plotter website (https://kmplot.com/analysis/). Patients were classified into high and low KAT8 expression subgroups based on the optimal cut-off value, defined as the value with the highest significance (lowest FDR). When multiple cut-off values yielded identical significance, the cut-off associated with the highest hazard ratio (HR) was selected for final analysis. Data were analyzed using GraphPad Prism 9 (GraphPad Software Inc) and presented as the mean ± standard deviation (SD). Statistical significance between two independent groups or among multiple groups was assessed using Student’s t-test and one-way ANOVA, respectively. A *P* value of < 0.05 was considered significant.

## Results

### Elevated KAT8 expression is correlated with poor prognosis in cervical cancer

To assess the clinical relevance of KAT8 in cervical carcinogenesis, its expression in cervical cancer was first examined. Immunohistochemical (IHC) analysis of tissue samples from 43 cervical cancer patients revealed significantly elevated KAT8 protein levels in tumor tissues compared to adjacent non-cancerous tissues (Figure 1A, B). Cellular-level investigations further supported these results. Western blot demonstrated increased KAT8 expression in four cervical cancer cell lines (C33A, Caski, SiHa, and HeLa) relative to normal cervical epithelial HcerEpic cells (Figure 1C). Notably, KAT8 expression is higher in HPV+ cervical cancer cell lines compared to the HPV-C33A cell line. The correlation between KAT8 expression and patient outcomes was then analyzed. Kaplan-Meier survival analysis revealed no significant association between elevated KAT8 expression and poor overall survival (OS) in cervical squamous cell carcinoma patients (*P =* 0.096) (Figure 1C). However, lower KAT8 expression correlated with significantly reduced recurrence-free survival (RFS) compared to higher expression (*P =* 0.014), suggesting a potential association with recurrence in cervical squamous cell carcinoma (Figure 1D). To evaluate the diagnostic potential of KAT8 expression, receiver operating characteristic (ROC) curve analysis was conducted. The area under the ROC curve (AUC) for KAT8 expression was 0.844 (CI = 0.671-1.000), demonstrating strong discriminatory ability between tumor and non-tumor tissues (Figure 1E). These results highlight the role of KAT8 in cervical cancer progression and its potential clinical diagnostic utility.

**Figure 1.**
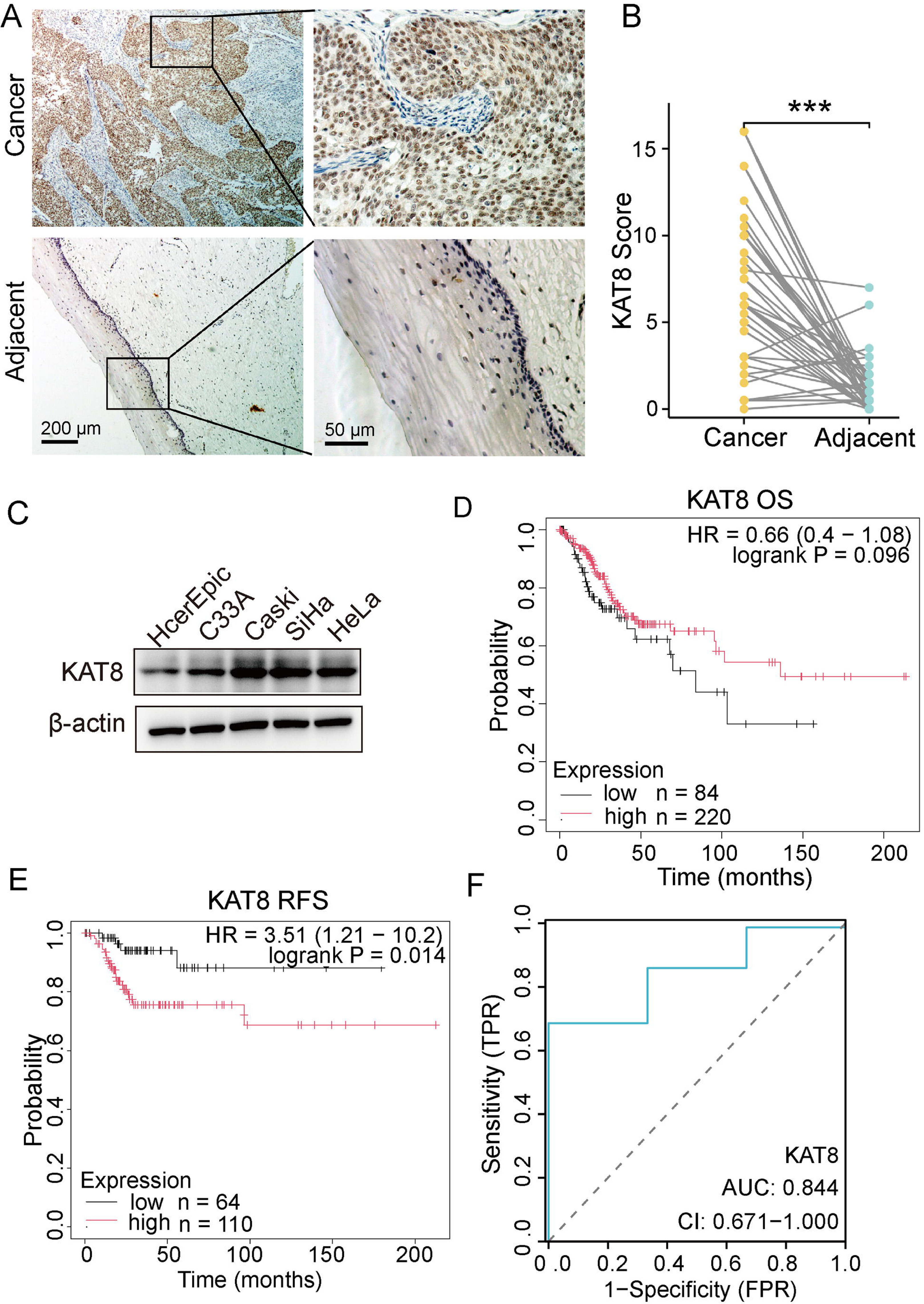
KAT8 expression in cervical cancer tissues and correlations to patient outcomes. A-B. IHC staining (A) and corresponding IHC scores (B) of KAT8 expression in cervical cancer tissues and adjacent normal tissues from 43 patients with cervical cancer (*** *P* < 0.001). C. Western blot detected the KAT expression in four cervical cancer cell lines (C33A, Caski, SiHa, and HeLa) compared to normal cervical epithelial HcerEpic cell. D. Kaplan-Meier survival analysis illustrating the association between KAT8 expression and OS. E. Patients with low KAT8 levels exhibited significantly longer RFS time than those with high KAT8 levels. F. ROC curve evaluating the diagnostic potential of KAT8 expression in distinguishing cervical cancer tissues from normal tissues.

### KAT8 promotes cervical cancer cell proliferation, migration, and invasion

To assess the impact of KAT8 on cervical cancer cell growth, KAT8 expression was silenced in HeLa (HPV18+) and SiHa (HPV16+) cervical cancer cell lines. Knockdown efficiency was verified by Western blot and RT-qPCR (Figure 2A, B). Both CCK-8 and colony formation assays confirmed that KAT8 silencing significantly impaired cell proliferation (Figure 2C-E). Further investigation into KAT8’s role in cell invasion revealed that shKAT8 cells exhibited markedly slower migration compared to control cells in wound healing assays (Figure 2F, G). Moreover, KAT8 knockdown led to a substantial reduction in both migratory and invasive capabilities (Figure 2H-J).

**Figure 2.**
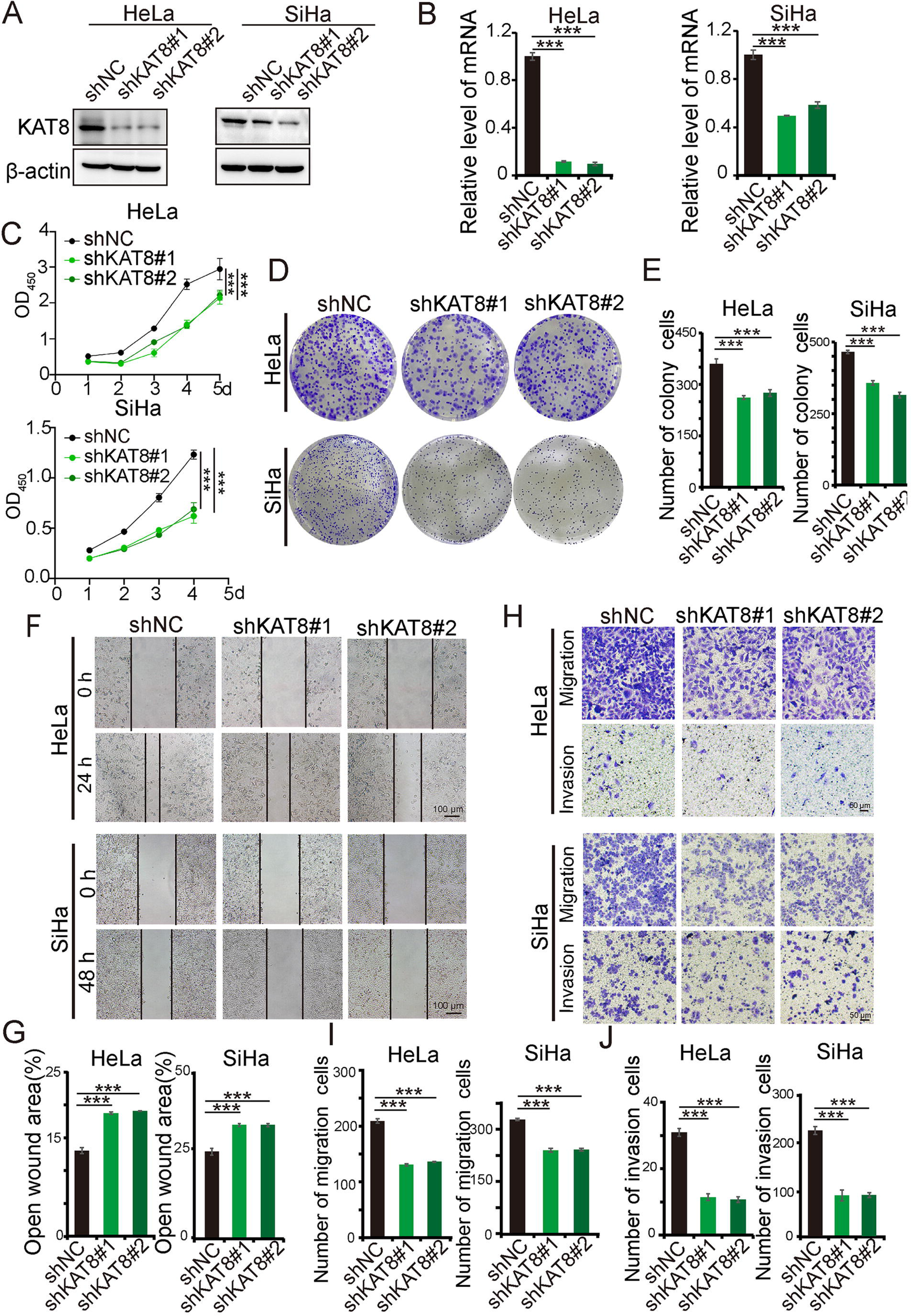
KAT8 knockdown suppresses the malignant biological behaviors of HPV16/18+ cervical cancer cells. A-B. Western blot (A) and RT-qPCR (B) analysis confirming KAT8 expression knockdown following lentiviral infection. C. CCK8 assay measuring cell viability in KAT8 knockdown (shKAT8) and control (shNC) cervical cancer cells. D-E. Colony formation assays assessing the effects of KAT8 knockdown on cervical cancer cell proliferation. F-G. Wound healing assays evaluating the impact of KAT8 inhibition on the motility of cervical cancer cells. H-J. Transwell migration and invasion assays analyzing the effects of KAT8 knockdown on cell migration and invasion capabilities. (*** *P* < 0.001).

KAT8-overexpressing cell lines (KAT8-OE) were subsequently generated in HeLa and SiHa cells (Figure 3A, B). KAT8 overexpression resulted in a notable increase in both cell proliferation (Figure 3C-F) and migration rate (Figure 3G, H) compared to control cells. Additionally, overexpression of KAT8 significantly enhanced the migratory and invasive potential of cervical cancer cells (Figure 3I-K). These results indicate that elevated KAT8 expression promotes malignant behaviors, including proliferation, migration, and invasion, in HPV16/18-positive cervical cancer cells.

**Figure 3.**
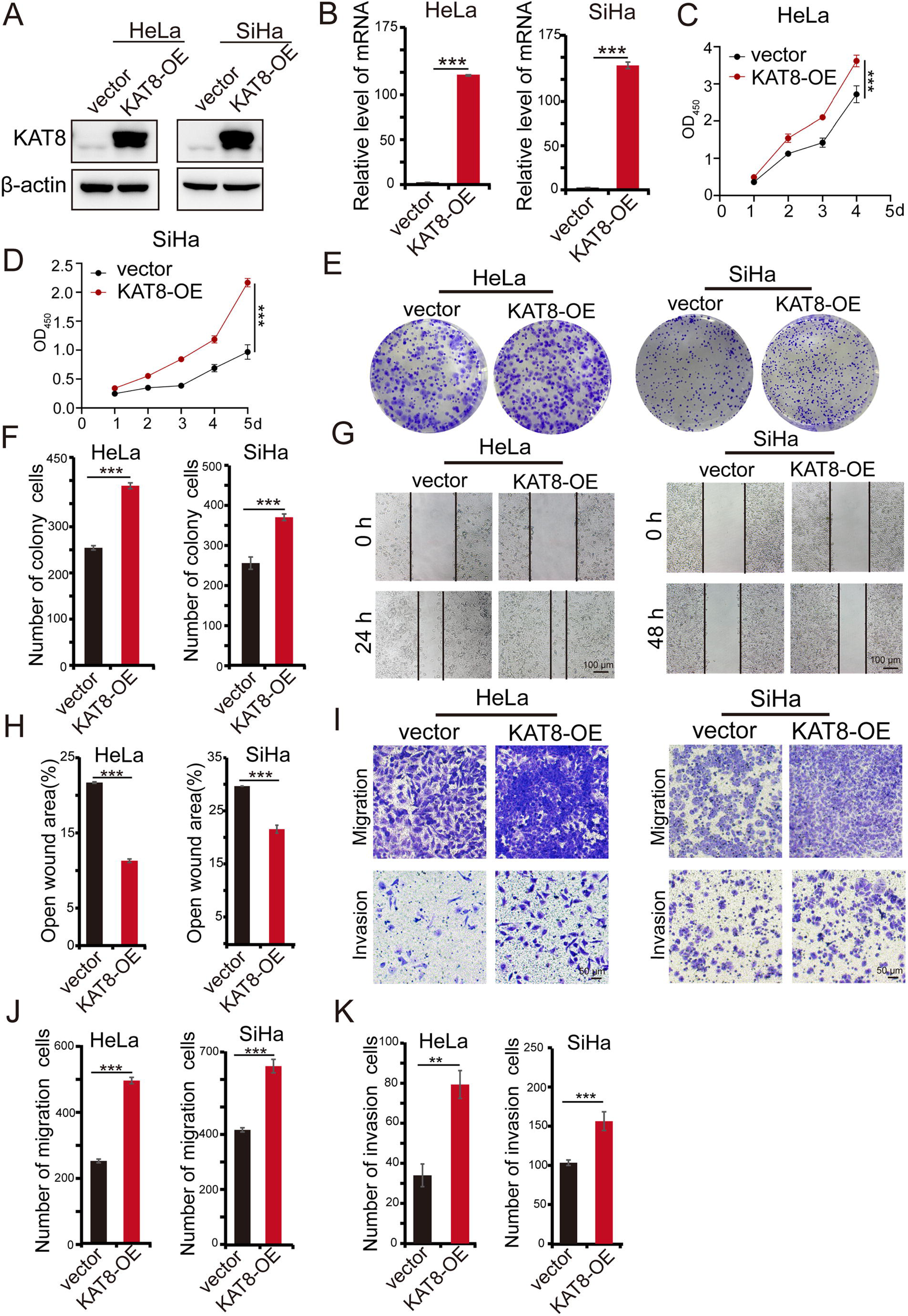
KAT8 promotes the growth and migration of HPV16/18+ cervical cancer cells. A-B. Protein (A) and mRNA (B) levels of KAT8 in cervical cancer cells transfected with vector or Flag-tagged KAT8 overexpression construct (KAT8-OE). C-D. CCK8 assay determining cell viability in cervical cancer cells transfected with vector or KAT8-OE. E-F. Colony formation assays examining the effect of KAT8 overexpression on cervical cancer cell proliferation. G-H. Wound healing assays evaluating the impact of KAT8 overexpression on cervical cancer cell motility. I-K. Transwell assays analyzing the capacity of migration and invasion in cervical cancer cells transfected with vector or KAT8-OE. (* *P* < 0.05, ** *P* < 0.01, *** *P* < 0.001).

### Low KAT8 expressions induce G1/S cell cycle arrest and DNA damage

To investigate the mechanisms by which KAT8 promotes proliferation in HPV16/18+ cervical cancer cells, its role in cell cycle progression and DNA damage repair was examined. Flow cytometry analysis revealed that KAT8 knockdown induced G1/S phase cell cycle arrest in both HeLa and SiHa cells (Figure 4A). Furthermore, TUNEL assay results indicated a substantial increase in apoptosis in HPV16/18+ cells following KAT8 knockdown (Figure 4B). FACS analysis also showed enhanced apoptosis levels in KAT8-knockdown cells (Figure 4C). To assess KAT8’s impact on DNA damage repair, the expression of the DNA damage marker γ-H2AX was measured. Elevated γ-H2AX levels in KAT8-knockdown HeLa and SiHa cells confirm that KAT8 depletion induce DNA damage in cervical cancer cells (Figure 4D). These data suggest that KAT8 is integral to maintaining chromatin stability in cervical cancer cells, although the precise regulatory mechanisms remain to be elucidated and warrant additional observation.

**Figure 4.**
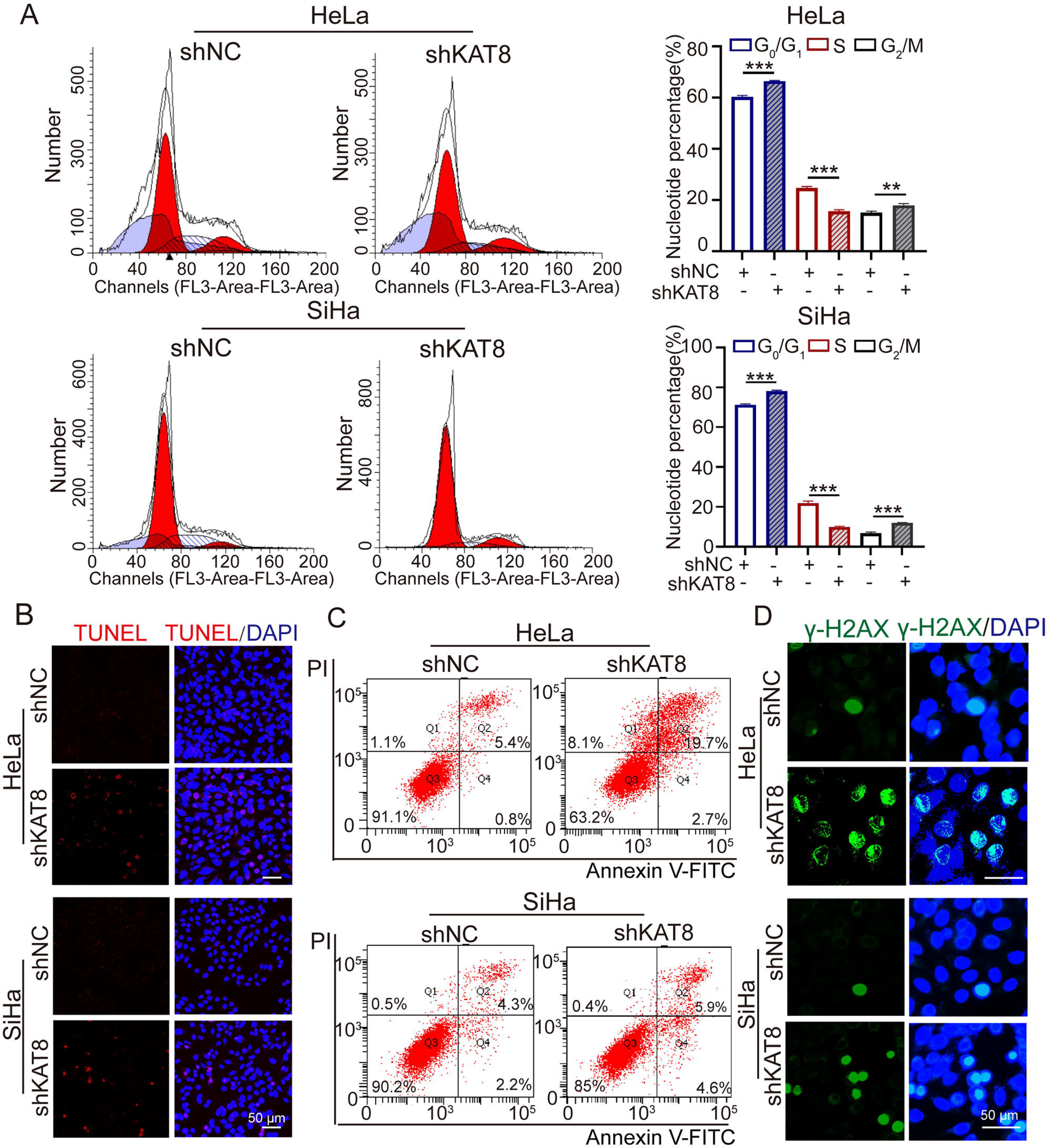
KAT8 affects G1/S phase transition and DNA damage repair in HPV16/18+ cervical cancer cell. Flow cytometry (FACS) analysis assessing the impact of KAT8 knockdown on the cell cycle distribution in HeLa and SiHa cells using PI staining. B. TUNEL assay was evaluated apoptosis cells in HeLa and SiHa cells with KAT8 knockdown. C. Cell apoptosis was evaluated with Annexin V/PI staining and FACS analysis in KAT8 knockdown HeLa and SiHa cells. D. Immunofluorescence staining for γ-H2AX in HeLa and SiHa cells, highlighting DNA damage response. (** *P* < 0.01, *** *P* < 0.001).

### KAT8 contributes to HPV infection and stimulate E7/pRb/E2F1 signaling axis

The role of KAT8 in cervical cancer was further investigated through RNA sequencing. A total of 555 differentially expressed genes (DEGs) were identified in both HeLa KAT8 and SiHa KAT8-OE cells. Of these, 139 genes exhibited consistent expression changes across both cell lines, including 43 upregulated and 96 downregulated genes (Figure 5A, B). GO enrichment analysis of the 139 DEGs revealed a strong association with processes related to cell proliferation and development, such as stem cell regulation. Notably, KEGG pathway enrichment analysis indicated that KAT8 was linked to HPV infection and cancer-related pathways (Figure 5C). These results establish a significant correlation between elevated KAT8 expression and HPV infection.

**Figure 5.**
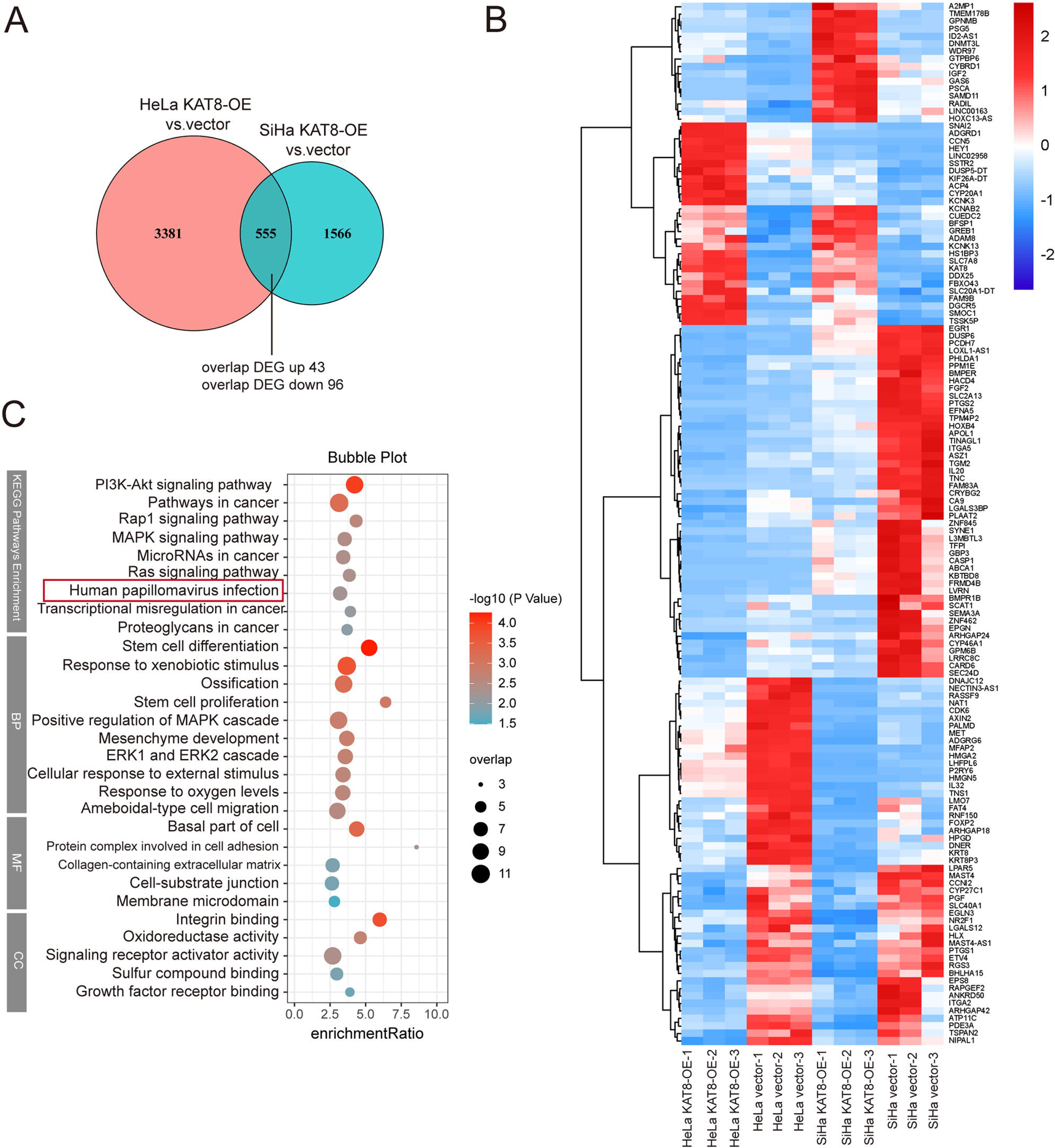
RNA-seq analysis of KAT8-OE and vector groups in cervical cancer cells. A. Venn diagram illustrating the overlap of differentially expressed genes (DEGs) in KAT8-OE cervical cancer cells compared to control cells. B. Heatmap depicting the expression profiles of overlapping DEGs with consistent changes between the KAT8-OE and vector groups. C. KEGG pathway analysis and GO enrichment analysis with 139 overlapping DEGs between the KAT8-OE and vector groups.

Cervical cancer is closely linked to HPV infection, prompting an investigation into the relationship between KAT8 and HPV. KAT8 protein expression was notably higher in HPV16/18+ cervical cancer patients compared to HPV-patients (Figure 6A, B). Among HPV-encoded proteins, the E7 oncoprotein plays a central role in tumorigenesis. E7 promotes cell cycle progression by inactivating the tumor suppressor pRb, thereby releasing E2F transcription factors from the pRb complex, advancing the cell cycle to the S phase, and driving uncontrolled cell proliferation [16]. To assess whether KAT8 regulates E7 expression, E7 protein levels were measured in cervical cancer cells after KAT8 interference. Results revealed that KAT8 overexpression increased E7 expression in HeLa and SiHa cells, while KAT8 knockdown reduced E7 protein levels, indicating a positive correlation between KAT8 and E7 expression (Figure 6C-D). Further examination of the E2F/pRb axis showed that KAT8 knockdown led to decreased E2F1 expression and increased pRb levels in cervical cancer cells (Figure 6C, D). Kaplan-Meier analysis revealed a significant positive correlation between KAT8 expression and E2F1 levels in cervical cancer tissues (r = 0.463, P < 0.001) (Figure 6F). To determine whether KAT8’s regulation of E7 oncogenic signaling was dependent on its acetyltransferase activity, HeLa cells were treated with MG149 to inhibit KAT8’s HAT activity, and E2F1/pRb/E7 protein expression was assessed by Western blot. The results in MG149-treated HeLa cells were consistent with those observed in KAT8 knockdown cells (Figure 6E), suggesting that KAT8’s acetylation activity is essential for the activation of E7-mediated oncogenic signaling. Furthermore, overexpression of E7 in KAT8 knockdown HeLa cells reversed the effects of KAT8 depletion on cell viability (Figure 6G-H). To further investigate the mechanism by which KAT8 regulated E7 expression, the transcriptional regulation of HPV18 E7 was examined. The long control region (LCR), located between the L1 and E6 open reading frames, consists of the proximal promoter and distal enhancer [17]. ChIP assays with anti-KAT8 antibodies demonstrated that KAT8 preferentially bound to the promoter regions over the enhancer regions (Figure 6I-J). Collectively, these data suggest that KAT8 promotes cervical cancer progression by modulating the E7/pRb/E2F1 signaling axis.

**Figure 6.**
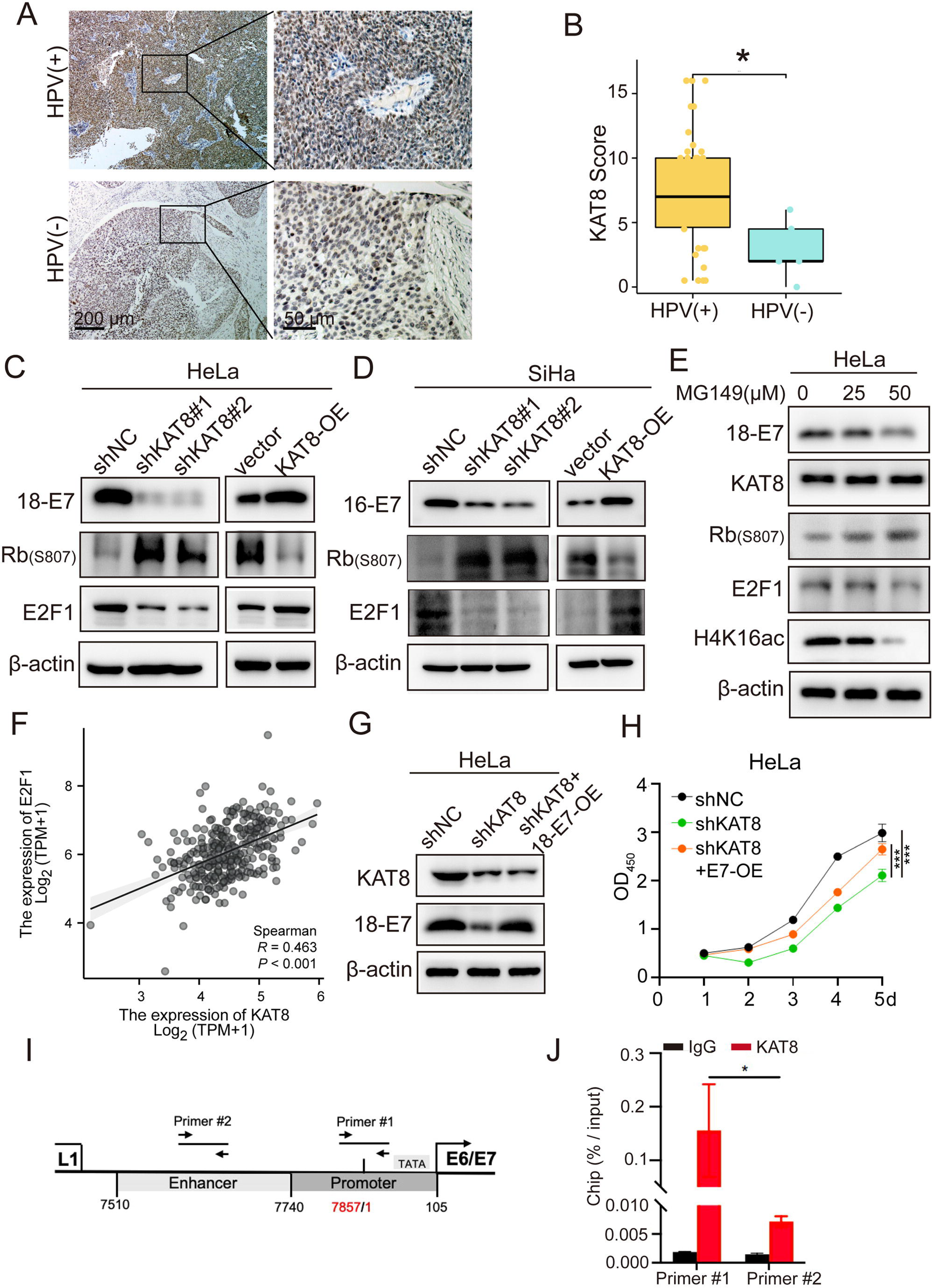
KAT8 promotes cervical cancer cell proliferation by stimulating E7/pRb/E2F1 axis. A-B. Representative IHC staining (A) and corresponding IHC scores (B) for KAT8 expression in HPV+ (n = 39) and HPV- (n = 5) cervical cancer samples. C-D. Western blot analysis of total protein extracts from HeLa (C) and SiHa (D) cells with KAT8 knockdown or overexpression, utilizing the same antibody panel. E. Western blot analysis of protein levels in HeLa cells treated with MG149 (25 µM and 50 µM) or vehicle (DMSO) for 24 hours. F. Kaplan-Meier analysis of the correlation between KAT8 and E2F1 expression in cervical cancer. G. Western blot analysis of 18-E7 protein expression in KAT8 knockdown HeLa cells transfected with HA-tagged E7, using an anti-E7 antibody. H. MTT assays evaluating the ability of E7 to rescue the inhibitory effect of KAT8 knockdown on HeLa cell proliferation. (*** *P* < 0.001). I. A schematic description of the HPV18 LCR. PCR fragments amplified in ChIP assays are indicated as primer #1 (promoter) and primer #2 (enhancer). J. The associations between KAT8 and HPV18 LCR were detected by ChIP assay performed with anti-KAT8 antibodies. (* *P* < 0.05)

### HPV16/17 E7 protein is acetylated by KAT8 in cervical cancer cells

KAT8, a key lysine acetyltransferase, regulates protein function through the acetylation of substrate proteins. To assess the impact of KAT8 on the acetylation of E7 and its role in cervical cancer progression, Flag-KAT8 and HA-HPV18-E7 (or HA-HPV16-E7) plasmids were co-transfected into HEK293T cells. Immunofluorescence (IF) staining revealed the colocalization of KAT8 and E7 in these cells (Figure 7A). Furthermore, co-immunoprecipitation (CoIP) experiments confirmed a direct interaction between KAT8 and HPV16/18-E7 (Figure 7B). To evaluate the acetylation of HPV16/18-E7, CoIP was performed to isolate exogenous HPV16/18-E7 from HEK293T cells. Western blot analysis using an anti-acetylated lysine antibody revealed significant acetylation of HPV16/18-E7 (Figure 7C). To determine whether KAT8 was responsible for E7 acetylation in cervical cancer cells, HeLa cells were treated with MG149 to inhibit KAT8’s HAT activity. Co-immunoprecipitation (CoIP) assays demonstrated that inhibiting MOF markedly reduced E7 acetylation (Figure 7D). Further investigations examined the functional impact of E7 acetylation on protein interactions, specifically between E7, pRb, and E2F1. To control for protein loading variations, pRb was used as a loading control in the total proteins immunoprecipitated with an anti-pRb antibody. As shown in Figure 7E, the interaction between pRb and E2F1 decreased, while the interaction between pRb and E7 increased in KAT8-OE HeLa cells. This suggests that KAT8 may enhance the interaction between E7 and pRb while diminishing the pRb/E2F1 interaction. These results indicate that KAT8-mediated acetylation of E7 may disrupt the pRb/E2F1 complex, leading to the release of E2F1 and promoting cervical cancer progression.

**Figure 7.**
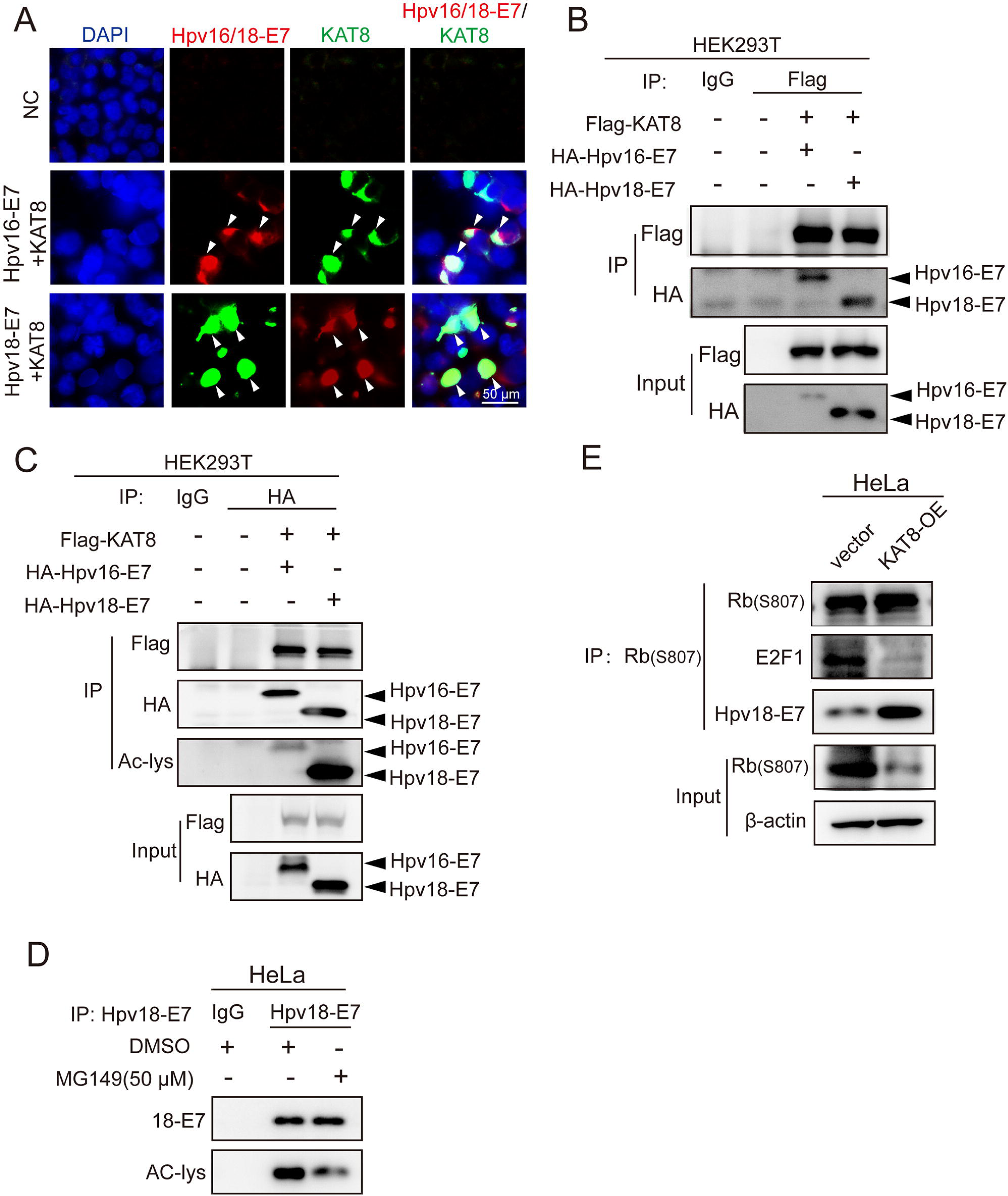
Acetylation of E7 protein and the effect of KAT8 on the E7/pRb/E2F1 interaction. A. Immunofluorescence staining showing co-localization of KAT8 (green) and E7 (red), indicated by yellow overlap. B. Co-immunoprecipitation assay using anti-Flag antibody to examine the interaction between KAT8 and E7 in HEK293T cells co-transfected with Flag-KAT8 and HA-HPV18-E7 (or HA-HPV16-E7) plasmids for 48 hours. C. Co-IP assay with anti-HA antibody confirming the acetylation of exogenous E7 protein, with acetylation levels assessed by Western blot using an anti-acetylated lysine antibody. D. Co-IP assay with anti-HPV18 E7 antibody to examine the E7 acetylation in HeLa cells treated with MG149 for 24 hours, followed by Western blot using an anti-acetylated lysine antibody. E. Immunoprecipitation analysis from KAT8-OE HeLa cells and control cells using anti-pRb antibody, followed by Western blot to examine the effect of KAT8 overexpression on the E7/pRb/E2F1 interaction.

## Conclusion

HPV-driven cervical cancer is associated with a favorable prognosis and higher incidence in younger patients. Despite the availability of effective vaccines, specific antiviral therapies for HPV and targeted treatments for HPV-associated cancers are still lacking. As a result, there is growing interest in the development of targeted therapies to enhance patient outcomes. HPV is a small, double-stranded DNA virus, with a genome that includes early genes (E1, E2, E4, E5, E6, and E7) responsible for viral gene expression, and late genes (L1 and L2) encoding capsid proteins. The E7 protein, in particular, plays a significant role in tumorigenesis, although the molecular mechanisms governing its function remain poorly understood.

KAT8, a histone acetyltransferase (HAT), is involved in numerous cellular processes, including gene transcription [11, 18, 19], DNA damage repair [20], apoptosis, and tumorigenesis[21]. It catalyzes the acetylation of H4K16 and partially acetylates H4K5 and H4K8, in addition to acetylating non-histone proteins such as p53 [22]. Dysregulation of KAT8 has been implicated in the initiation and progression of several cancers. However, its direct impact on tumorigenesis and the underlying regulatory mechanisms are unclear. Early studies have shown that KAT8 knockdown in HeLa cells results in the downregulation of oncogenes like HOXA9 [23], suggesting its potential oncogenic role. Furthermore, KAT8 deletion exacerbates the genotoxic effects of cisplatin in HeLa cells, and mitochondrial KAT8 has been shown to repair respiratory defects in these cells [15]. Our data corroborate these findings, demonstrating that KAT8 is upregulated in cervical cancer tissues and significantly associated with recurrence-free survival (RFS), but not with overall survival (OS). Given the heterogeneity of cervical cancer, further investigation with a larger sample size is required to explore potential correlations between KAT8 expression and other clinicopathological factors, such as tumor stage, grade, and lymph node involvement. Such studies would provide a more comprehensive understanding of KAT8’s clinical relevance in cervical cancer. Additionally, KAT8 knockdown suppresses the proliferation, migration, and invasion of cervical cancer cells in vitro. Studies have demonstrated that KAT8 knockout induces DNA damage, G2/M cell cycle arrest, and genomic instability in embryonic fibroblasts [20, 24]. However, the role of KAT8 in cervical cancer cells remains unexplored. In this study, KAT8 knockdown in HeLa and SiHa cells resulted in enhanced apoptosis, G1/S cell cycle arrest, and DNA damage, offering new insights into the potential function of KAT8 in HPV-associated cancers.

The HPV oncoprotein E7 disrupts the G1 checkpoint and induces genomic instability, both of which are integral to cervical carcinogenesis. Previous studies have shown that pan-HDAC inhibitors influence HPV-18 DNA amplification and cellular DNA synthesis [25]. These results suggest a potential regulatory role for HATs in cervical cancer cell biology; however, the specific function of these enzymes remains insufficiently explored. This study establishes that E7 is essential for KAT8-mediated promotion of cervical cancer cell proliferation, implicating KAT8 in the progression of HPV-driven cervical cancer.

E2F1, a key transcription factor, regulates multiple cellular processes, including proliferation, DNA repair, apoptosis, and cell cycle progression. It activates genes required for the G1/S phase transition. Dysregulation of E2F1 expression has been implicated in various cancers, including cervical carcinoma [26], with its overexpression contributing to cervical cancer progression. HPV E7 promotes the degradation of pRb, freeing E2F1 from the pRb/E2F1 complex and driving malignant transformation of the cervical epithelium [27, 28]. While pRb is a known target of E7, E2F1 is a critical downstream effector of pRb. While pRb is a well-established target of E7, E2F1 serves as a key downstream effector of pRb. In this study, KAT8 knockdown resulted in a marked reduction of E2F1 expression, establishing a positive correlation between KAT8 levels and E2F1 activity. These results suggest that KAT8 modulates the G1/S transition in E7-expressing cells through regulation of E2F1.

Post-translational modifications play a fundamental role in modulating protein activity [29]. A significant finding in this study was the identification of an interaction between KAT8 and E7, along with the acetylation of the E7 protein. Although direct evidence for KAT8-mediated acetylation of endogenous E7 remains absent, the data suggest that KAT8 is essential for the interaction between E7 and the pRb/E2F1 axis in cervical cancer cells.

In summary, KAT8, a histone acetyltransferase, is upregulated in cervical cancer and plays a critical role in the proliferation of HPV-positive cervical cancer cells. Silencing KAT8 in HPV16/18+ cervical cancer cells can markedly impair oncogenic phenotypes *in vitro*. Mechanistically, KAT8 drives cervical cancer progression by modulating the E7/pRb/E2F1 signaling axis. These results advance the understanding of the molecular mechanisms underlying cervical cancer progression and propose KAT8 as a potential therapeutic target for HPV-associated malignancies. However, the current study is limited by its preliminary nature, necessitating further investigation, including *in vivo* studies, to fully elucidate the role of KAT8 in cervical cancer progression and its therapeutic potential.

---

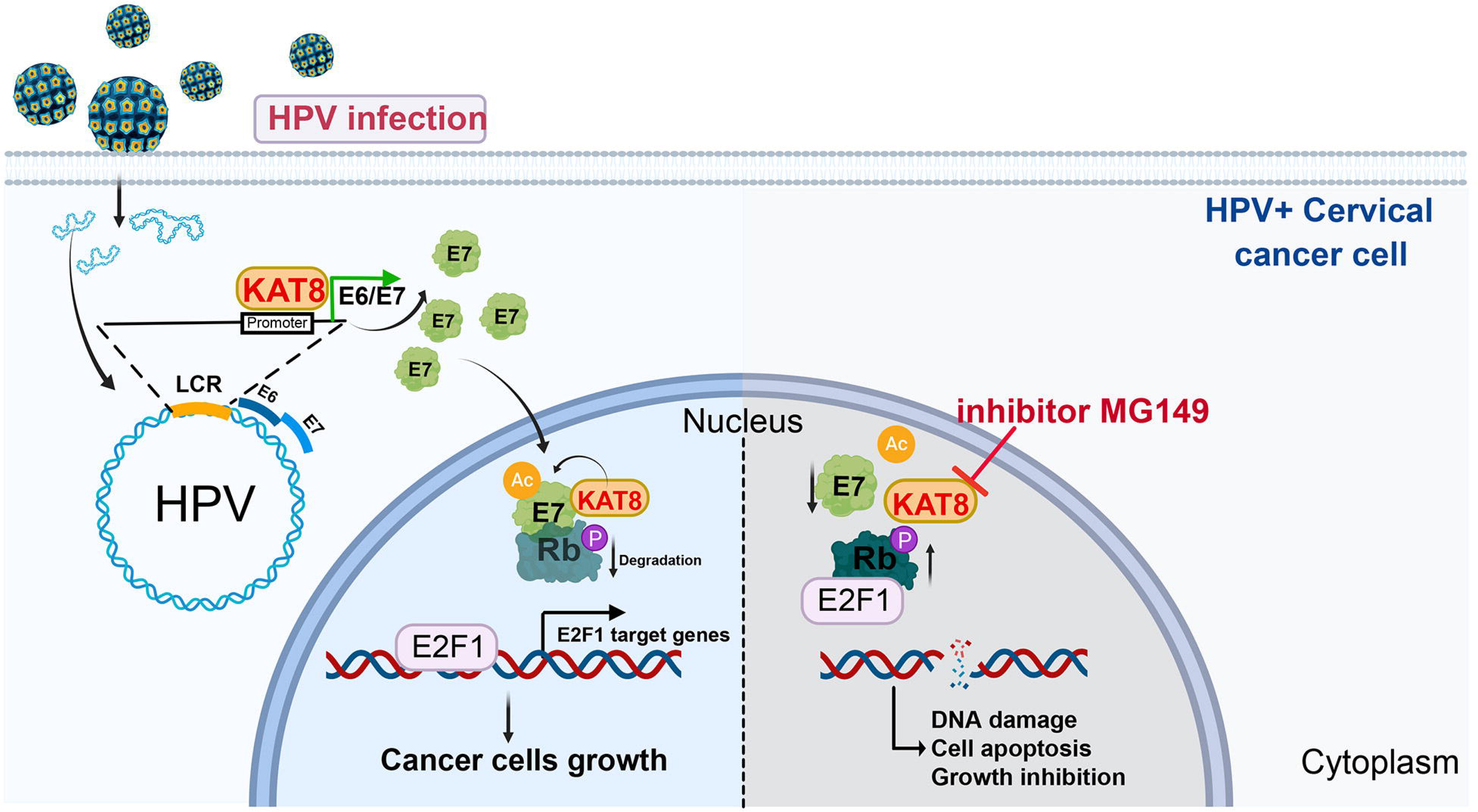

